# Nutritional stress-induced regulation of microtubule organization and mRNP transport by HDAC1 controlled α-tubulin acetylation

**DOI:** 10.1101/2023.02.09.527826

**Authors:** Frank Wippich, Vaishali, Marco L. Hennrich, Anne Ephrussi

**Author notes:** Correspondence should be addressed to A.E. Contributed equally (Frank Wippich, Vaishali).

## Abstract

In response to nutritional stress, microtubules in cells of the *Drosophila* female germline are depleted from the cytoplasm and accumulate cortically. This triggers aggregation of mRNPs into large reticulated sponge bodies and oogenesis arrest. Here, we show that hyperacetylation of α-tubulin at lysine 40 (K40) alters microtubule dynamics and sponge body formation. We found that depletion of histone deacetylase 1 (HDAC1) by RNAi phenocopies the nutritional stress response, causing hyperacetylation of α-tubulin, as well as accumulation of maternally deposited mRNPs in reticulated sponge bodies. Using *in vitro* and *in vivo* studies, we identify HDAC1 as a direct regulator of α-tubulin K40 acetylation status. In well-fed flies, HDAC1 maintains low levels of α-tubulin acetylation, enabling the microtubule dynamics required for mRNP transport. Using quantitative phosphoproteomics we identify nutritional stress-induced changes in protein phosphorylation that act upstream of α-tubulin acetylation, including phosphorylation of HDAC1 at S391, which reduces its ability to deacetylate α-tubulin. These results reveal that *Drosophila* HDAC1 senses and relays the nutritional status, which regulates germline development through modulation of cytoskeleton dynamics.

## Introduction

*Drosophila* oogenesis is tightly regulated by nutrient availability, which is sensed by the mother and transmitted to the developing germline cells via insulin signalling (Burn et al. 2015). Upon nutrient deprivation, oogenesis arrests and egg chambers form large reticulated mRNP aggregates, termed sponge bodies, in both the oocyte and interconnected nurse cells (Wilsch-Bräuninger et al. 1997; Snee and Macdonald 2009). Given similarities in protein composition, sponge bodies have also been referred to as processing bodies (Decker and Parker 2006; Shimada et al. 2011), but are distinct morphologically and in their association with ER-like cisternae (Wilsch-Bräuninger et al. 1997; Snee and Macdonald 2009). Formation of sponge bodies during nutritional stress depends in part on loss of intact microtubules, as microtubule depolymerizing drugs, such as colchicine, induce sponge body formation (Shimada et al. 2011), although the exact mechanism remains unclear. A systemic insulin signal has been shown to activate insulin/TOR signaling in follicle cells, which regulates both microtubule organization and sponge body formation in the underlying germline cells, and reduced insulin signaling in nutrient deprived conditions leads to cytoplasmic depletion and cortical condensation of microtubules (Burn *et al*, 2015). However, neither the receiving nor the processing factors in the germline target cells have been identified thus far.

Post-translational modifications of α-tubulin are involved in microtubule dynamics and stability (Janke and Montagnac 2017). Acetylation of α-tubulin at K40, carried out primarily by α-tubulin acetyltransferase 1 (αTAT1), is known to alter microtubule dynamics by enhancing the flexibility of microtubules, thereby protecting them from mechanical stresses and making them more long-lived (Akella *et al*, 2010; Kalebic *et al*, 2013; Portran *et al*, 2017). So far, changes in α-tubulin acetylation have been shown to affect multiple cellular processes, including intracellular trafficking(Reed et al. 2006), and nutrient deprivation has been shown to induce hyperacetylation of α-tubulin (Geeraert et al. 2010).

Histone deacetylases (HDACs), have been shown to regulate mRNP dynamics during stress and to coordinate stress granule formation via microtubule-based transport mechanisms in mammalian cells (Kwon et al. 2007). Moreover, cytoplasmic HDACs, most prominently class II HDACs, such as cytoplasmic HDAC6, can deacetylate α-tubulin *in vitro* and *in vivo* (Hubbert et al. 2002; Matsuyama et al. 2002; Zhang et al. 2003).

Here, we identified deacetylation of α-tubulin on K40 by *Drosophila* HDAC1 as a regulatory step in the nutritional stress response in the female germline. We found that depletion of HDAC1 by RNAi phenocopies the nutritional stress response, causing hyperacetylation of α-tubulin, as well as accumulation of maternally deposited mRNPs in reticulated sponge bodies. HDACs are themselves regulated by post-translational modifications, such as phosphorylation, to either modulate enzymatic activity, dimerization or subcellular localization (Segré and Chiocca 2011; Yang and Seto 2008; Khan et al. 2013). Using quantitative phosphoproteomics of well-fed and nutritionally deprived egg chambers, we show that HDAC1 is hyperphosphorylated on S391 upon nutritional stress. We further demonstrate that phosphorylation of S391 plays a role in regulating the deacetylase activity of HDAC1 specifically towards α-tubulin. Taken together these data indicate that HDAC1 is required to transmit nutritional information by modulating microtubule dynamics and microtubule-based mRNA transport in the *Drosophila* female germline.

## Results

### α-Tubulin acetylation affects oocyte mRNA transport

Upon nutrient deprivation, egg chambers form large reticulated sponge bodies that contain maternal mRNAs, including *oskar* mRNA (Burn et al. 2015; Wilsch-Bräuninger et al. 1997; Snee and Macdonald 2009) (Fig. 1A). Formation of sponge bodies is promoted by insulin signaling-mediated alterations in microtubule dynamics and organization, and specifically the depletion of cytoplasmic microtubules in egg chambers (Burn et al. 2015; Shimada et al. 2011). We found that α-tubulin in *Drosophila* ovarian lysates is hyperacetylated at K40 upon nutritional stress, provoked by withdrawal of the standard protein-rich fly diet (Fig. 1B). While nutritional stress results in the depletion of the cytoplasmic pool of microtubules, an accumulation of microtubules has been observed in the cortical region of nurse cells (Shimada et al. 2011). Here, we found that the cortically enriched microtubules are hyperacetylated in nutrient-deprived egg chambers (Fig. 1C and Fig. S1). Thus, hyperacetylation of α-tubulin at K40 correlates with the enrichment of cortical microtubules in nurse cells and the formation of sponge bodies.

**Figure 1:**
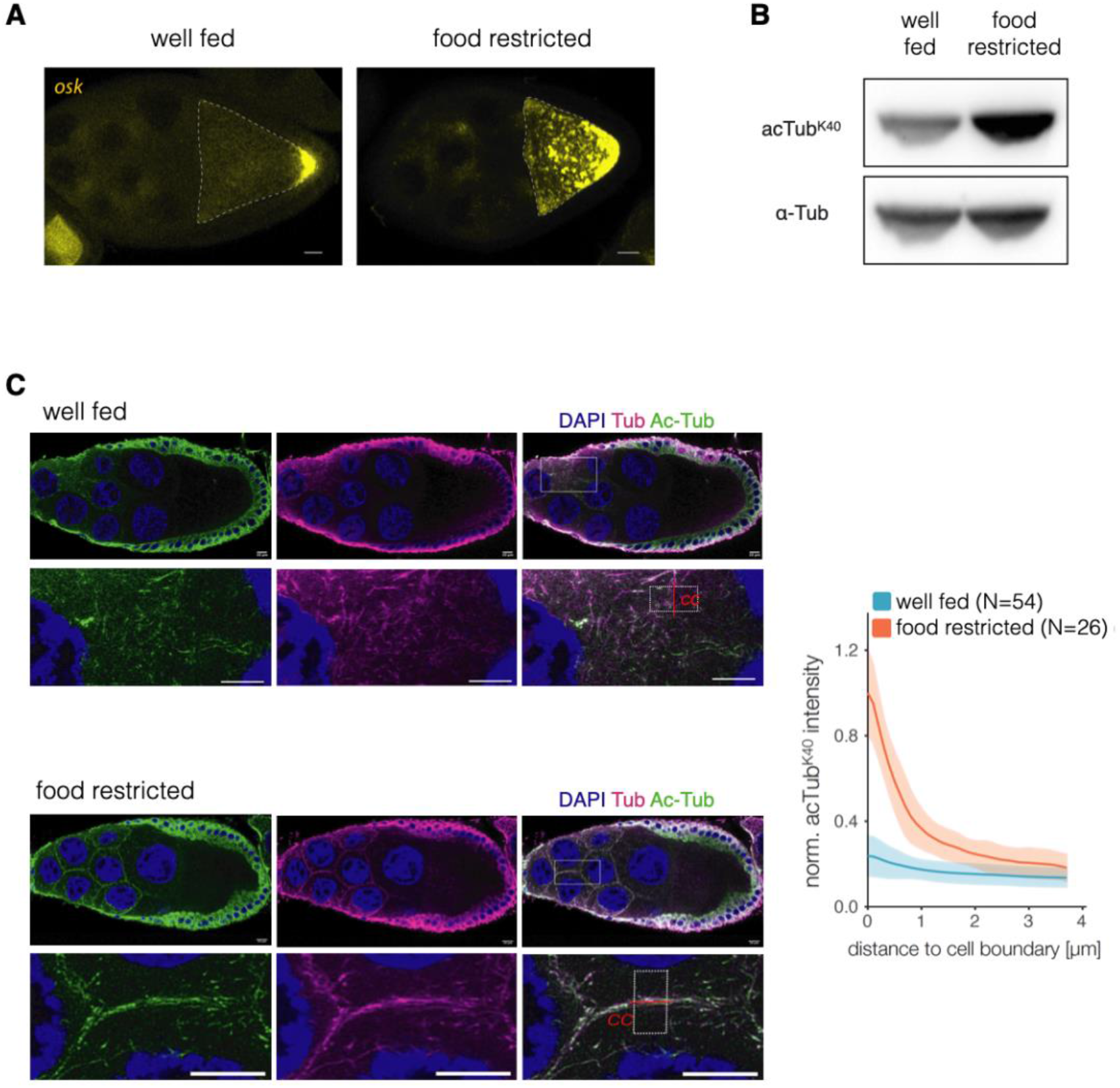
α-Tubulin acetylation affects mRNA transport in the developing oocyte. (A) *oskar* mRNA localization in egg chambers from well-fed and nutritional deprived flies visualized by smFISH. The oocyte is highlighted by a dashed line. (B) Western blot analyse of α-tubulin K40 acetylation in ovarian lysates from well-fed and nutritional deprived flies. (C) Acetylation of α-tubulin in egg chambers from well-fed and nutritionally deprived flies stained for tubulin (tub), acetylated α-tubulin K40 (Ac-Tub) and DAPI. Lower panels show a magnification of the rectangular area highlighted in the upper panels on the right (merged signals). Graph shows a quantification of the fluorescence signal at cell to cell boundaries (cc, red line) between nurse cells in ovaries of well fed (N = 54) and food restricted (N = 26) flies. The fluorescent profile of a squared area (within white dashed line) parallel to the cell boundary was used for quantification, and fluorescence intensity was normalised to the maximum observed intensity. Shaded areas represent SD. Scale bar: 10μm

To determine the role of α-tubulin acetylation at K40, we made use of transgenic flies that overexpress either wild type, acetylation deficient (K40R) or acetylation mimetic (K40Q) α-tubulin (Bhattacharjee 2012). At mid oogenesis, *oskar* mRNA localizes to the posterior of the oocyte. Overexpression of either wild type or acetylation deficient K40R α-tubulin did not affect *oskar* mRNA localization (Fig. 2A,B) whereas the acetylation mimetic mutant α-tubulin K40Q did, resulting in substantial *oskar* mRNA accumulation throughout the ooplasm, with only a smaller proportion being localized to the posterior pole (Fig. 2C). The *oskar* mRNA levels were similar for the three lines, as quantified by qRT-PCR (Fig. S2). The dominant effect of α-tubulin K40Q suggests that changes in α-tubulin dynamics caused by mimicking hyperacetylation are involved in the accumulation of *oskar* mRNPs into sponge bodies, as seen during nutritional stress.

**Figure 2:**
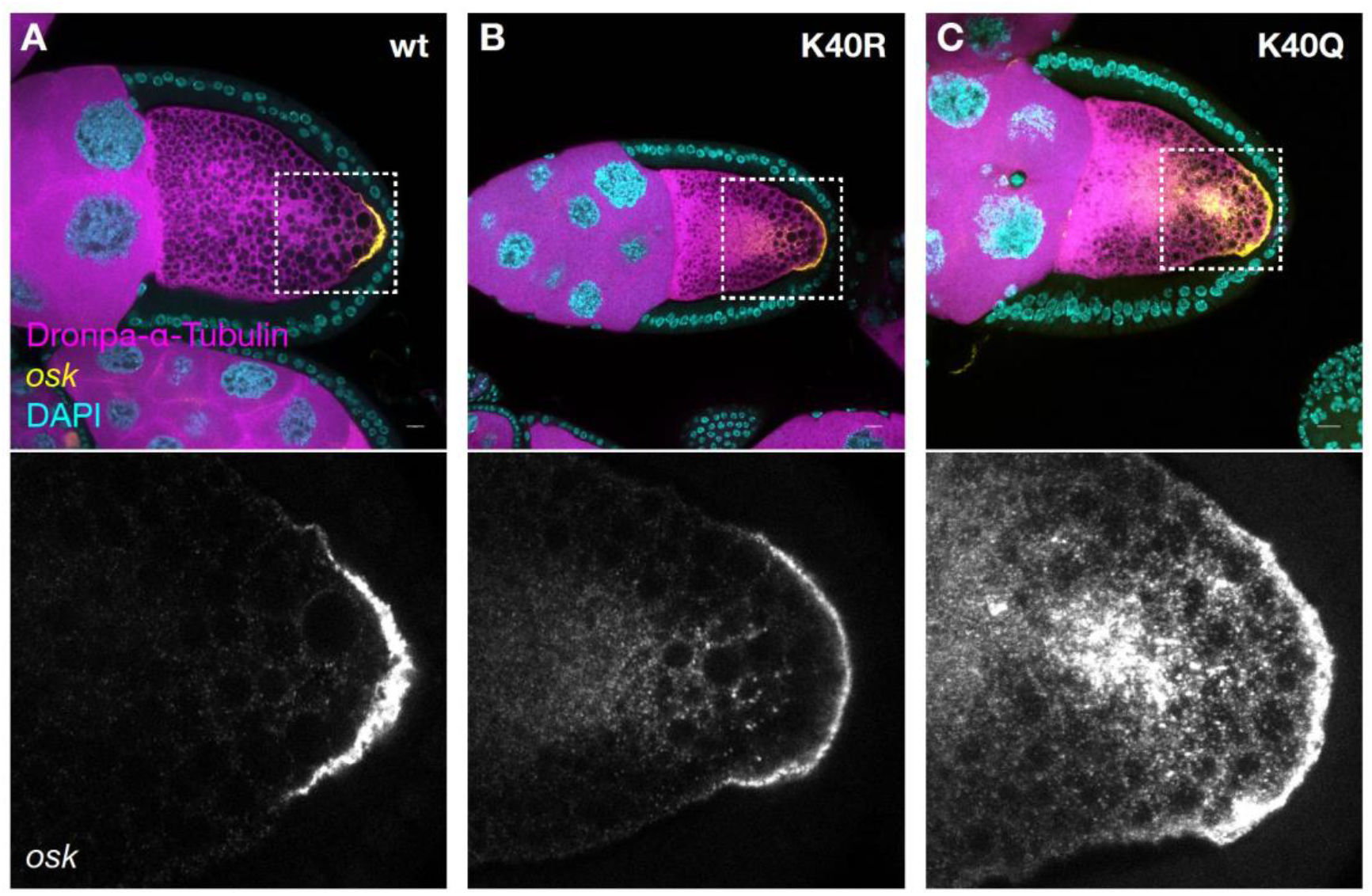
Acetylation mimetic α-tubulin K40Q affects the localization of *oskar* mRNA. (A-C) Representative confocal images of egg chambers expressing Dronpa-α-tubulin wild-type, K40R and K40Q mutants. *oskar* mRNA localization was visualized by smFISH. Scale bar: 10 μm

### HDAC1 modulates microtubule dynamics and mRNA transport

Given cytoplasmic HDACs can deacetylate α-tubulin *in vitro* and *in vivo* (Hubbert et al. 2002; Matsuyama et al. 2002; Zhang et al. 2003) we assayed a panel of *Drosophila* class 1, 2a, and 2b HDACs for their impact on α-tubulin acetylation under well-fed conditions (Fig. 3A). We found that germline-specific knock down of HDAC1 using two different RNAi lines substantially increased α-tubulin K40 acetylation (Fig. 3B), as seen during nutritional stress (Fig. 1B). Knock down of other HDACs, including HDAC6, which has been shown to be the main mammalian α-tubulin deacetylase (Hubbert et al. 2002; Matsuyama et al. 2002), showed only minor effects on α-tubulin K40 acetylation levels in the *Drosophila* female germline.

**Figure 3:**
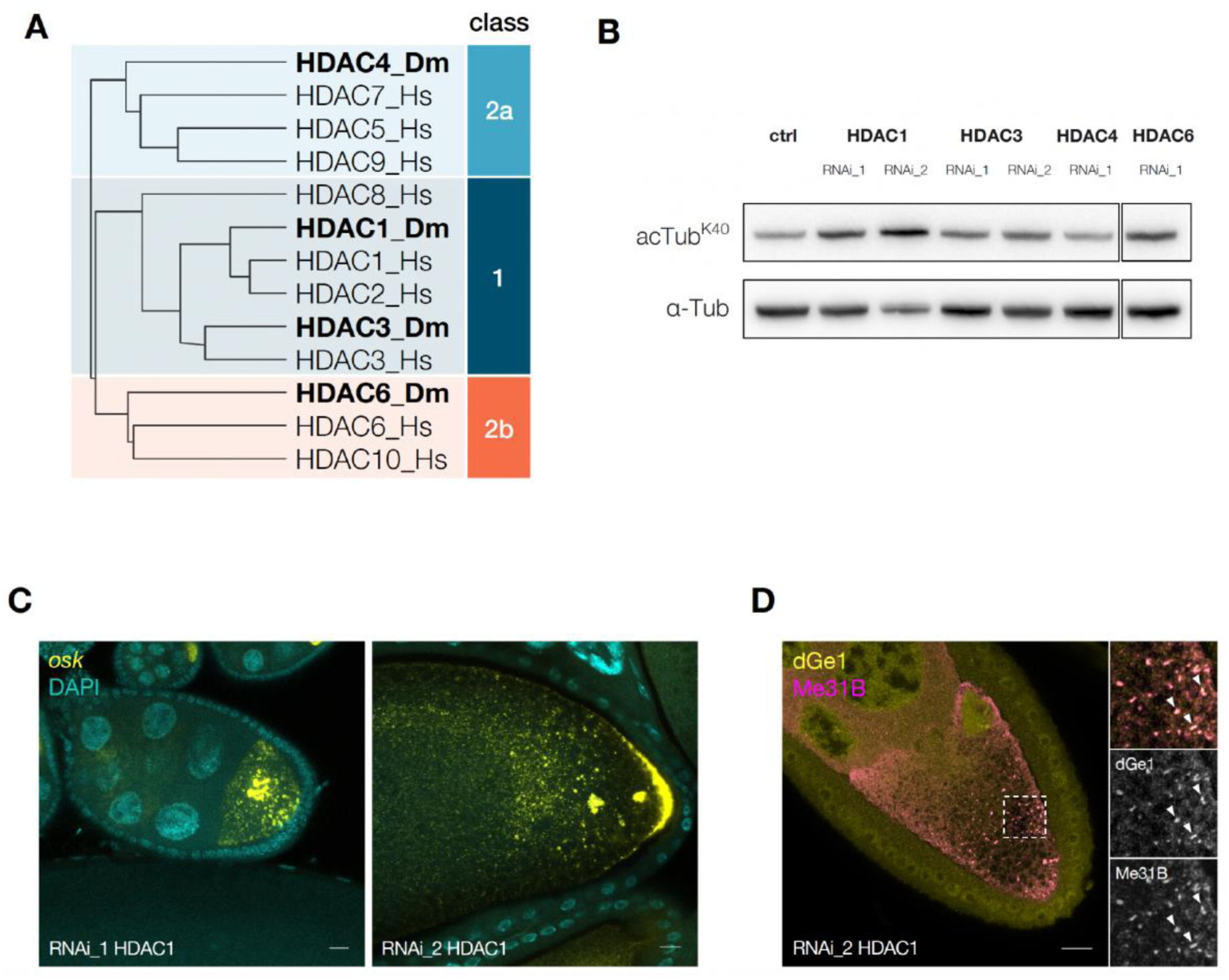
HDAC1 RNAi affects α-Tubulin acetylation and mRNP transport. (A) Phylogenetic analysis of class 1, 2a, and 2b HDAC proteins from Homo sapiens (Hs) and *Drosophila melanogaster* (Dm) using Clustal Omega. (UniprotIDs for Hs: Q13547, Q92769, O15379, Q9UQL6, Q9UBN7, Q8WUI4, Q9BY41, Q9UKV0, Q969S8; for Dm: Q94517, Q7KTS4, Q9VYF3, Q86NK9) (B) Western blot analysis of α-tubulin K40 acetylation from ovaries in which HDAC1, HDAC3, HDAC4 and HDAC6 were knocked down using RNAi in the germline. (C) *oskar* mRNA localization visualized using smFISH in HDAC1 knock-down egg chambers using two different RNAi lines. (d) Immuno-fluorescence of dGe1 and Me31B accumulating in sponge bodies in an HDAC1 knock-down egg chamber. Scale bar: 10μm

We also found that *oskar* mRNA transport was strongly perturbed upon HDAC1 knock down, as evident from the aggregation of *oskar* mRNA in large puncta, resembling sponge bodies, in the oocyte cytoplasm (Fig. 3C). Indeed, when we investigated the subcellular localization of the mRNA granule marker proteins Me31B and dGe1, we found that despite the lack of nutritional stress, both proteins aggregated in sponge bodies (Fig. 3D). Furthermore, knockdown of HDAC3, HDAC4 and HDAC6 did not lead to aggregation of *oskar* mRNPs under well fed conditions, indicating the phenotype is specific to HDAC1 (Fig. S3). This suggests that HDAC1 maintains low levels of α-tubulin acetylation in the *Drosophila* female germline under well fed conditions, presumably promoting the microtubule dynamics required for normal mRNP transport.

### *Drosophila* HDAC1 directly deacetylates α-Tubulin

To test whether *Drosophila* HDAC1 directly deacetylates α-tubulin we produced recombinant HDAC1 protein using a baculovirus-mediated expression system in insect cells. When incubated with tubulin purified from porcine brain, we found that HDAC1 was able to reduce the acetylated α-tubulin level in a concentration dependent manner, showing that *Drosophila* HDAC1 can efficiently deacetylate α-tubulin (Fig. 4A). Overexpression of HDAC6 has been shown to reduce the amount of acetylated α-tubulin in human cells, while other HDACs showed no effect on α-tubulin acetylation levels (Hubbert et al. 2002; Matsuyama et al. 2002). We found that overexpression of GFP-tagged *Drosophila* HDAC1 in S2R+ cells resulted in substantially reduced acetylation of α-tubulin K40 in transfected compared to untransfected cells (Fig. 4B and Supplemental Fig. S4A). This indicates that, in contrast to human HDAC1 (Hubbert et al. 2002; Matsuyama et al. 2002), *Drosophila* HDAC1 directly deacetylates α-tubulin, both *in vitro* and *in vivo*.

**Figure 4:**
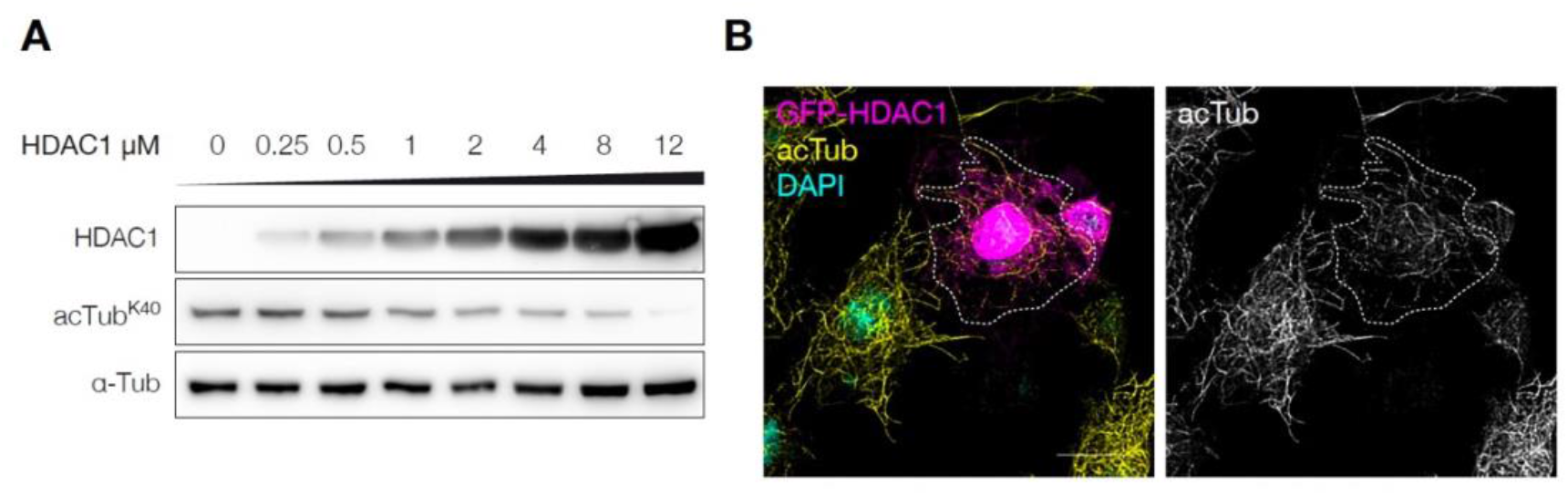
HDAC1 deacetylates α-tubulin *in vitro* and *in vivo*. (A) *In vitro* deacetylation assay using indicated amounts of recombinant HDAC1 with 1 μM porcine tubulin, incubated for 2 hours at 25°C and analysed by western blot. (B) S2R+ cells transiently transfected with GFP-HDAC1, stained 24 hours post transfection for acetylated α-tubulin K40 and DAPI. A transfected cell is highlighted by the dashed line. Scale bar: 10 μm

### Phosphorylation-mediated control of microtubule acetylation

Insulin/TOR signalling has been shown to transmit information about nutritional status to *Drosophila* follicle cells and to regulate the starvation response (Burn et al. 2015). However, it is unclear which signalling pathways mediate the nutritional stress response within the germline. Supplementing *ex vivo* cultured egg chambers with insulin suppresses the formation of sponge bodies (Shimada et al. 2011). We therefore wondered if sponge body formation is sensitive to changes in the phosphorylation status of proteins, given phosphorylation underlies the signaling cascade events of the insulin/TOR pathway. When cultured for 2.5 hours *ex vivo* without insulin (Fig. 5A), egg chambers showed a typical starvation response, with *oskar* mRNA accumulating in sponge bodies (Fig. 5B), as is seen *in vivo* (Fig. 1A). However, the addition of a broad-spectrum phosphatase inhibitor (PhosSTOP) substantially reduced the formation of *oskar* mRNA containing sponge bodies (Fig. 5B), suggesting that phosphorylation is required for the nutritional stress response. Moreover, hyperacetylation of α-tubulin caused by *ex vivo* culturing of egg chambers without insulin could also be suppressed by the addition of the phosphatase inhibitor (Fig. 5C). Taken together, this suggests that phosphorylation events acting upstream of α-tubulin acetylation/deacetylation regulate microtubule dynamics and subsequent formation of sponge bodies in response to nutrition.

**Figure 5:**
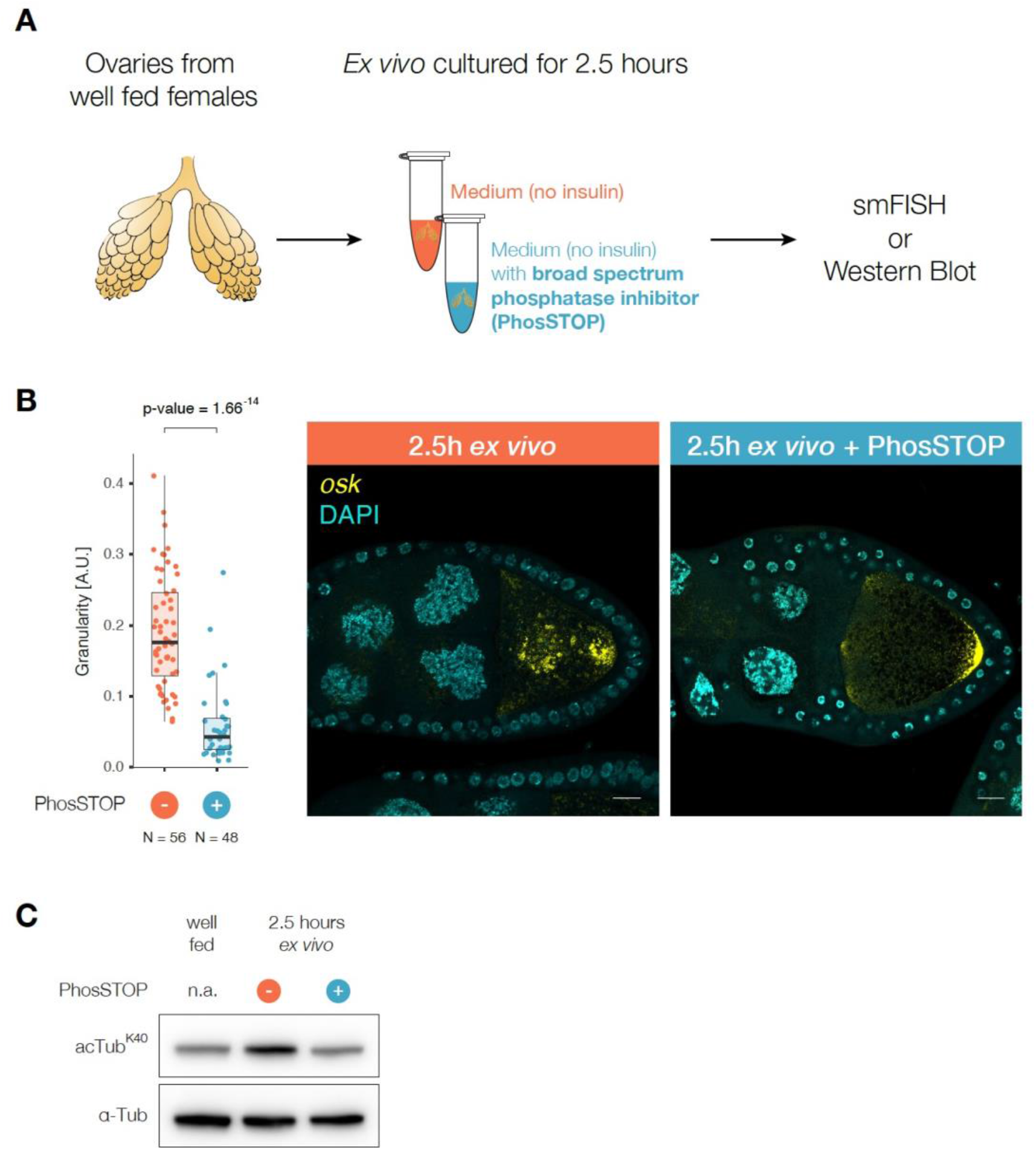
Phosphorylation events act upstream of sponge body formation and α-tubulin acetylation. (A) Schematic representation of *ex vivo* culturing assay of ovaries. Ovaries from well-fed females were dissected manually and transferred to medium or medium supplemented with a broad spectrum phosphatase inhibitor (PhosSTOP). After an incubation for 2.5 hours, ovaries were subjected to staining using smFISH or analysis by western blot. (B) smFISH staining for *oskar* mRNA in egg chambers cultured with or without PhosSTOP for 2.5 hours *ex vivo*. Statistical significance was calculated using Welch Two Sample t-test. (C) Acetylation of α-tubulin K40 from ovaries cultured with or without PhosSTOP for 2.5 hours *ex vivo* was analysed by western blot. Scale bar: 10μm

### Phosphorylation dynamics upon nutritional stress

In order to identify the signalling events involved in the nutritional stress response *in vivo*, we analysed the phosphorylation state of proteins from well-fed and nutrient deprived egg chambers. TiO_2_ enrichment of phospho-peptides followed by quantitative mass-spectrometry was used to quantitatively monitor 16975 phosphorylation sites in 2822 proteins (Supplemental Table S1). We found that the phosphorylation-sites of insulin/TOR signalling-related proteins were significantly less phosphorylated upon nutritional stress (Fig. 6A,B and Supplemental Fig. Table S1) consistent with somatic insulin and TOR signalling having been shown to transmit information about the nutritional status to the germline and to regulate the starvation response (Burn et al. 2015).

**Figure 6:**
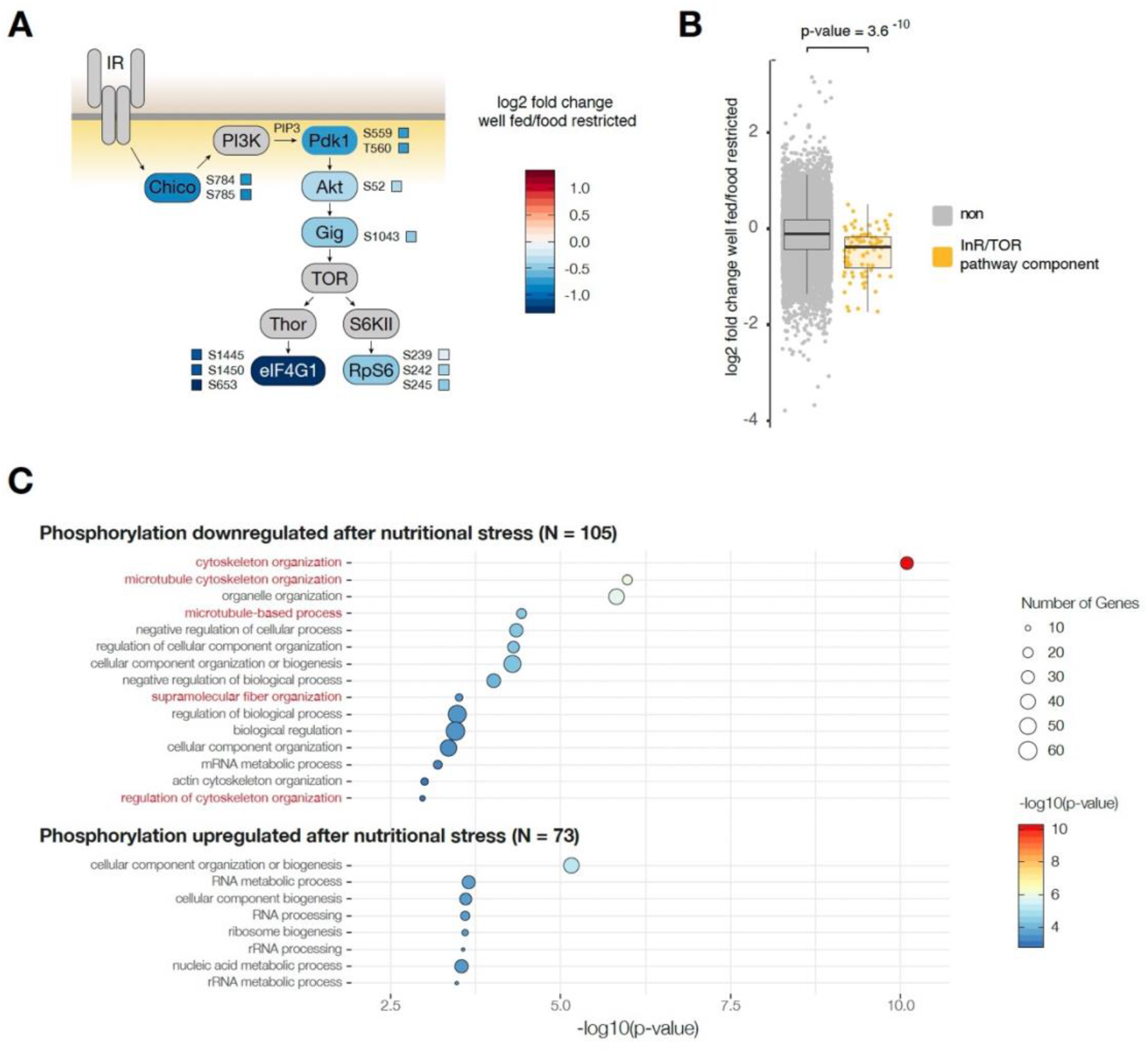
Nutritional stress induced changes in the phosphoproteome of *Drosophila* egg chambers. (A) Schematic representation of the insulin/TOR signalling pathway. Individual phosphorylation sites of relevant proteins which were analysed by quantitative phosphoproteomics are highlighted. (B) Quantification of all phosphorylation sites of proteins in the insulin/TOR signalling pathway. Statistical significance was calculated using Welch Two Sample t-test. (C) GO-term enrichment analysis of proteins whose phosphorylation sites were significantly down- or upregulated in response to nutritional stress (Benjamini-Hochberg corrected).

We asked which biological and molecular functions are associated with proteins whose phosphorylation states were altered upon nutritional stress. Proteins whose phosphorylation sites were significantly downregulated (p-value < 0.05) showed an enrichment of gene ontology (GO) terms associated with cytoskeleton organization, microtubule cytoskeleton organization, microtubule-based processes, supramolecular fiber organization, and regulation of cytoskeleton organization (Fig. 6C and Supplemental Table S2). Proteins whose phosphorylation sites were significantly upregulated showed no significant enrichment of GO terms related to the cytoskeleton. Thus, post-translational phosphorylation modifications play a role in transmitting nutritional status information to regulate cytoskeletal dynamics in the *Drosophila* female germline.

### HDAC1 activity during nutritional stress is regulated by phosphorylation

HDACs are themselves regulated by post-translational modifications, such as phosphorylation, to either modulate enzymatic activity, dimerization or subcellular localization (Segré and Chiocca 2011; Yang and Seto 2008; Khan et al. 2013). We identified seven phosphorylation sites in HDAC1 (Fig. 7A), of which only Serine 391 (S391) showed a significant increase in phosphorylation in response to nutritional stress compared to the remaining 6 phosphorylation sites, which were not significantly altered. S391 is located in a disordered region C-terminal to the histone deacetylase catalytic domain and the residue is highly conserved from fly to mouse, rat and human (Fig. 7B). In humans the corresponding sites in HDAC1 (S393) and HDAC2 (S394) have been shown to be phosphorylated (Segré and Chiocca 2011). Phosphorylation of S421/S423 in mammalian HDAC1 enzymes regulates catalytic activity (Loponte et al. 2016; Pflum et al. 2001). The effect of phosphorylation at S393 and at sites other than S421/S423 in mammalian HDAC1 is presently unclear.

**Figure 7:**
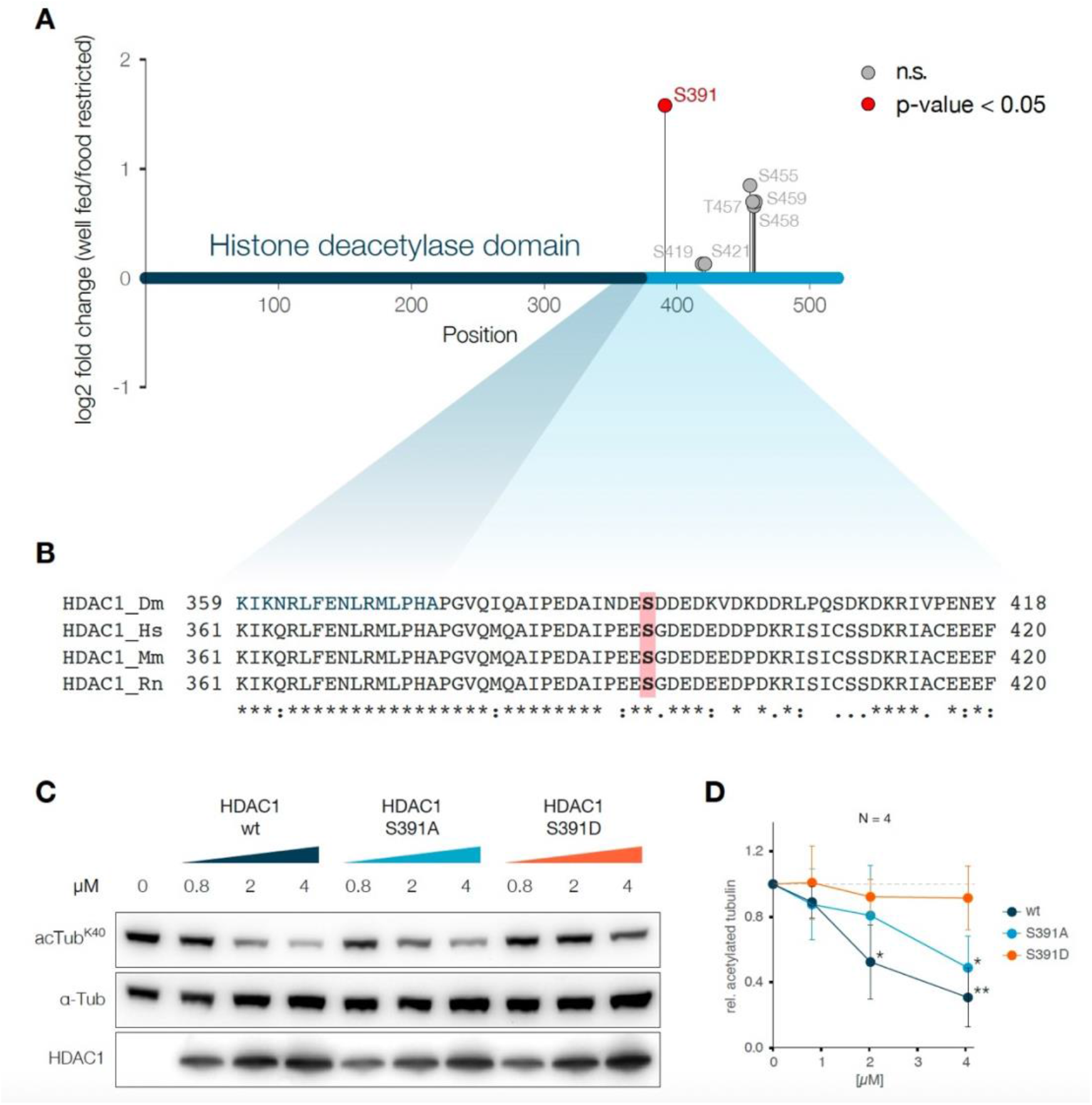
HDAC1 phosphorylation and the effect on α-tubulin deacetylation activity. (A) Visualization of *Drosophila* HDAC1 protein and the phosphorylation sites identified by quantitative phosphoproteomics. (B) Alignment of the conserved region around S391 of HDAC1 from *Drosophila melanogaster* (Dm), Homo sapiens (Hs), Mus musculus (Mm) and Rattus norvegicus (Rn). (C) Western blot analysis of *in vitro* deacetylation assay using recombinant wild type, mutant S391A and S391D HDAC1 at indicated concentrations and purified tubulin (1 μM), incubated for 1 hour at 25°C. (D) Quantification of western blot analysis of *in vitro* deacetylation assays as in C) (N = 4). Error bars represent SD. Statistical significance was calculated using Welch Two Sample t-test: * p-Value <0.05, ** p-Value < 0.01.

To study the effect of S391 phosphorylation in *Drosophila* HDAC1, we produced phosphorylation deficient (S391A) and phosphorylation mimetic (S319D) HDAC1 proteins using a baculoviral expression system. We found that phosphorylation deficient HDAC1 S391A had a deacetylation activity comparable to wild type HDAC1 on purified tubulin (Fig. 7C,D). The phospho-mimetic HDAC1 S391D failed to deacetylate α-tubulin K40, suggesting that, in *Drosophila*, phosphorylation of S391 suppresses the enzymatic activity of HDAC1 towards α-tubulin.

Consistent with the *in vitro* data, wild type and GFP-HDAC1-S391A overexpression in S2R+ cells significantly reduced the level of acetylated α-tubulin K40 in transfected cells compared to non-transfected neighbouring cells (Supplemental Fig. S4A). In contrast, phospho-mimetic GFP-HDAC1-S391D was unable to reduce the level of acetylated α-tubulin, consistent with the diminished activity of phospho-mimetic HDAC1 S391D *in vivo*. Phosphorylation has also been shown to regulate nucleocytoplasmic trafficking of HDACs, which might influence acetylation dynamics due to reduced substrate availability (Yang and Seto 2008). However, neither the phospho-mimetic (S391D) nor the phospho-deletion (S391A) mutation altered the nuclear to cytoplasmic ratio of GFP-HDAC1 in cultured cells (Supplemental Fig. S4B). Taken together, our data show that HDAC1 is dephosphorylated at S391 in response to nutritional stress, which decreases its ability to deacetylate α-tubulin.

Surprisingly, we found no substantial difference between wild-type HDAC1, HDAC1 S391A, and HDAC1 S391D in an assay that used short artificial peptide substrates to measure deacetylase activity (Supplemental Fig. S5A). This suggests that rather than directly influencing the enzymatic activity of the deacetylase domain, phosphorylation at S391 regulates substrate recognition or release. The disordered C-terminus of HDAC1 and its phosphorylation state may regulate substrate recognition, an idea consistent with a recent report suggesting a role for the unstructured N-terminus of mammalian HDAC6 in α-tubulin recognition (Ustinova et al. 2020). We found substantial differences in the amount of copurified α-tubulin when preparing the different phospho-mutant HDAC1s from insect cells (Supplemental Fig. S5B). Compared to wild type HDAC1, HDAC1 S391A copurified less and HDAC1 S391D much more α-tubulin, suggesting that the phosphorylation status of S391 plays a role in regulating the binding and/or release of the α-tubulin substrate.

## Discussion

Microtubule dynamics are critical for mRNA transport and mRNP granule formation in the *Drosophila* female germline (Burn et al. 2015; Zimyanin et al. 2008). During nutritional stress α-tubulin and microtubules are depleted from the cytoplasm and accumulate cortically. The loss of cytoplasmic microtubules may result in the aggregation of mRNPs and the formation of reticulated sponge bodies.

Here, we show that hyperacetylation of α-tubulin K40 correlates with altered microtubule dynamics and the formation of sponge bodies during nutritional stress in the *Drosophila* female germline (Fig. 1A,B). A similar mechanism operates in mammalian cells, where nutritional stress has been shown to induce the hyperacetylation of α-tubulin, a phenomenon crucial for the formation of autophagosomes and starvation-induced autophagy (Geeraert et al. 2010). Since the starvation response in egg chambers also leads to the induction of autophagy (Barth, Szabad et al. 2011), it is possible that microtubule hyperacetylation promotes recruitment of autophagy inducing factors during the starvation response. Additionally, it is possible that in mammalian cells, as in *Drosophila* egg chambers, changes in microtubule dynamics contribute to mRNP aggregation and stress granule formation upon nutrient deprivation.

Further, we show that acetylation mimetic α-tubulin K40Q can phenocopy the effects of nutritional stress (Fig. 2C). Acetylation of α-tubulin at K40 is known to alter microtubule dynamics by enhancing the flexibility of microtubules, thereby protecting them from mechanical stresses and making them more long-lived (Portran et al. 2017). Thus, we propose a model in which nutritional stress-induced hyperacetylation of α-tubulin alters microtubule organization, depleting them from the cytoplasmic pool of dynamic microtubules, which then leads to the aggregation of mRNPs (Fig. 8).

**Figure 8:**
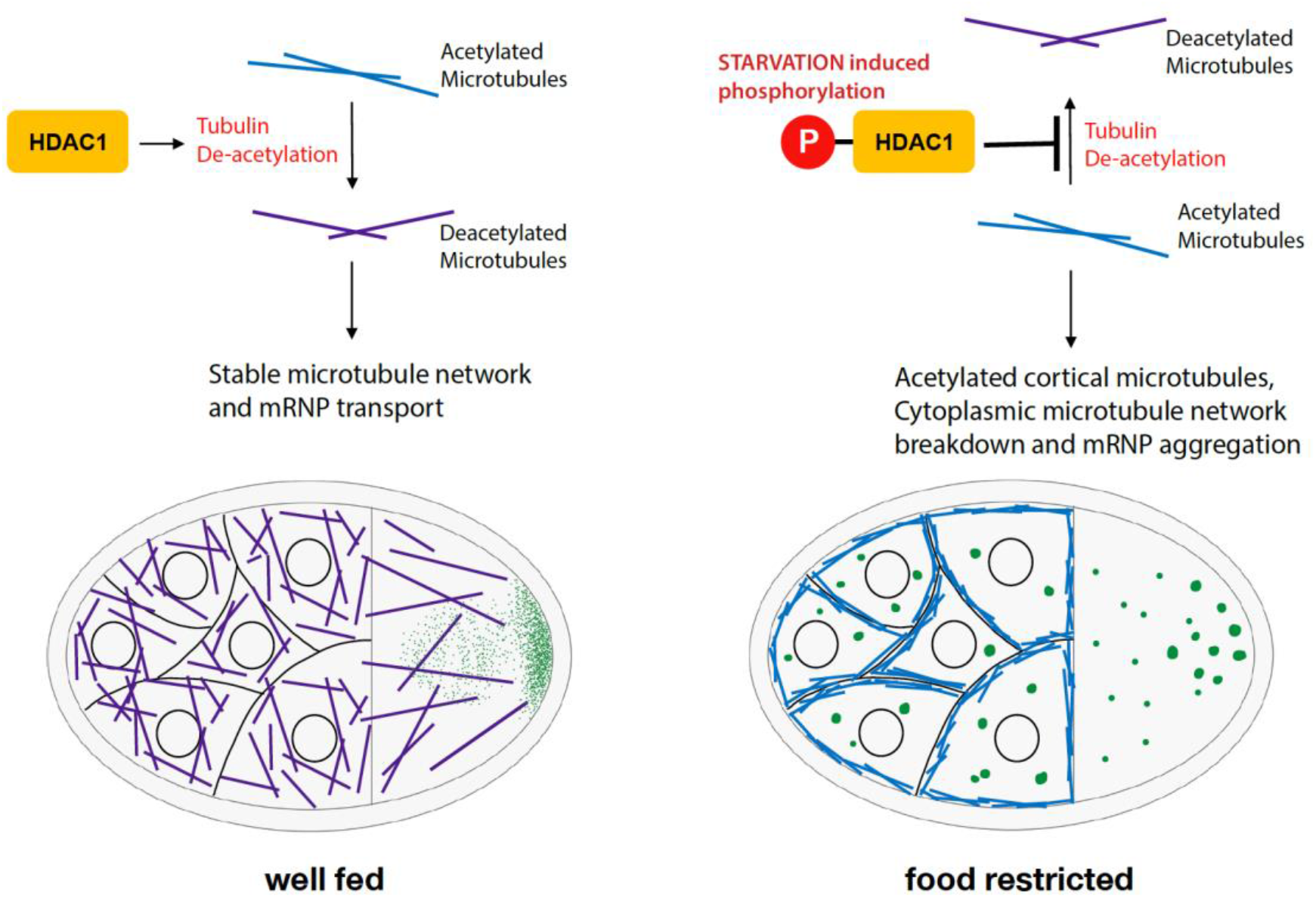
Model for regulation of starvation response by HDAC1: In well fed flies, HDAC1 maintains low levels of acetylated microtubules in the germline. Upon food deprivation, phosphorylation of HDAC1 reduces its ability to deacetylate microtubules, resulting in an accumulation of acetylated microtubules near the cell cortex.

Class II HDACs, such as the cytoplasmic HDAC6, have been shown to deacetylate α-tubulin (Hubbert et al. 2002; Matsuyama et al. 2002; Zhang et al. 2003). Here, we identified class I HDAC1 as the deacetylase principally responsible for the deacetylation of α-tubulin in the *Drosophila* female germline (Fig. 3B). We further show that *Drosophila* HDAC1, in contrast to mammalian HDAC1 (Hubbert et al. 2002), directly deacetylates α-tubulin *in vitro* and *in vivo* (Fig. 4A,B). Why HDAC1 is more involved in the regulation of α-tubulin K40 acetylation than HDAC6 in the *Drosophila* female germline remains to be clarified. *Drosophila* HDAC1 and HDAC6 have anticorrelated temporal expression patterns during developmental stages: HDAC1 is higher expressed in the early developmental stages whereas HDAC6 expression is highest in the adult fly (Cho et al. 2005). Therefore, specific HDACs might carry out similar functions and have overlapping substrates throughout developmental stages.

Phosphorylation-mediated regulation of HDACs has been shown to affect enzymatic activity, subcellular localization and dimerization (Segré and Chiocca 2011; Yang and Seto 2008; Khan et al. 2013). Quantitative phosphoproteomics revealed that HDAC1 S391 is significantly phosphorylated in response nutritional stress (Fig. 7A), and the phospho-mimetic mutant HDAC1 S391D shows substantially reduced α-tubulin K40 deacetylation activity *in vitro* and *in vivo* compared to wild type or phospho-deletion S391A HDAC1 (Fig. 7C,D and Supplemental Fig. S4A). However, the phospho-mimetic mutant HDAC1 S391D could deacetylate a short peptide substrate as well as the wild type enzyme or the phosphorylation deficient mutant HDAC1 S391A (Supplemental Fig. S5A). This suggests a regulatory mechanism which involves the phosphorylation of S391 for substrate recognition and/or release. Indeed, removal of the N-terminal microtubule binding domain of HDAC6 has been shown to have no effect on the deacetylase activity towards short peptides, but was crucial for the recognition of α-tubulin (Ustinova et al. 2020). We found substantial differences in the amount of copurified α-tubulin when preparing the different phospho-mutant HDAC1s from insect cells (Supplemental Fig. S5B). Compared to wild type HDAC1, HDAC1 S391A copurified less and HDAC1 S391D much more α-tubulin, suggesting that the phosphorylation status of S391 within the disordered C-terminus plays a role in regulating the binding and/or release of the α-tubulin substrate.

Protein kinase CK2, and PKA and Aurora kinases have been shown to phosphorylate human class I HDACs (Tsai and Seto 2002; Loponte et al. 2016). When analysing our phospho-proteomics data with respect to known kinase consensus motifs, we found a significant increase in phosphorylation of CK2 consensus motifs upon nutritional stress (Supplemental Fig. S6A). Furthermore, S391 of *Drosophila* HDAC1 is embedded in a predicted target of the CK2 consensus sequence (X-(S/T)-X-X-(D/E), whose corresponding site in human HDAC1 has been shown to be phosphorylated by CK2 *in vitro* (Bian et al. 2013). CK2 has been shown to play important roles in carbohydrate metabolism, by regulating insulin production and secretion, as well as regulating the activity of enzymes involved in carbohydrate storage and metabolism in hormone sensitive cells (Al Quobaili & Montenarh, 2012). Hence, it was of interest to test if, in *Drosophila* egg chambers, the starvation response is also dependent on CK2 mediated phosphorylation. However, culturing ovaries *ex vivo* in the presence of the selective CK2 inhibitor tetrabromocinnamic acid (TBCA) did not affect the nutritional stress-induced hyperacetylation of α-tubulin (Supplemental Fig. S6B). At very high concentrations, CK2 inhibition even increased the hyperacetylation of α-tubulin K40. Thus, we found no evidence that CK2 is involved in the nutritional stress induced phosphorylation of HDAC1 and the subsequent hyperacetylation of α-tubulin in the *Drosophila* female germline.

Taken together, our results reveal that *Drosophila* HDAC1 plays a role in nutritional status sensing, which regulates germline development. Changes in microtubule dynamics are likely controlled by a number of different signalling pathways. We found several examples of microtubule associated proteins whose phosphorylation status was significantly altered in response to nutritional stress, such as Tau, Futsch and its interaction partner Ank2, Spectrin, mei-38 (Supplemental Fig. S7). We also found regulators of microtubule dynamics, such as Ensconsin, Stathmin, BicC and Tral, and other cytoskeleton associated proteins, such as hts, to be less phosphorylated. For several of these (e.g. Tau, Ensconsin and Stathmin), phosphorylation has been shown to have a function in the regulation of binding to microtubules (Nouar et al. 2016; Stoothoff and Johnson 2005; Sung et al. 2008).

## Materials and methods

### Fly stocks

*w*^*1118*^ was used as wild type. Oregon-R (Bloomington Stock 5#) was used for mass extraction of ovaries for phosphoproteomics. RNAi against HDAC1 (RNAi_1: Bloomington Stock #36800; RNAi_2: Bloomington Stock #33725), HDAC3 (RNAi_1: Bloomington Stock #64476; RNAi_2: Bloomington Stock #34778), HDAC4 (RNAi_1: Bloomington Stock #34774; RNAi_2: Bloomington Stock #28549), HDAC6 (RNAi_1: Bloomington Stock #34072) and control RNAi (Bloomington Stock #35573) were driven by MTD-Gal4 (Bloomington Stock # 31777). K40R and K40Q α-tubulin expressing lines were from (Bhattacharjee 2012).

### Antibodies and reagents

The following antibodies were used: Mouse anti-α-Tubulin (Sigma, T5168), mouse anti-acetylated-α-Tubulin (Sigma, T7451), mouse anti-Me31B (Nakamura et al. 2001), rat anti-dGe1 (Fan et al. 2011), rabbit anti-GFP antibody-Alexa Fluor 488 conjugate (Invitrogen, A-21311) and rabbit anti-HDAC1 (gift of Jürg Müller).

### Fluorescent in situ hybridization

Ovaries from 2-day-old females, well-fed or protein deprived, were dissected into 2% v/v PFA (Electron Microscopy Sciences, #15710), 0.05% v/v Triton X-100 (Sigma, P1379) in PBS (pH 7.4) and were fixed for 20 minutes. After two washes with PBSTX (0.1% Triton x-100 in PBS), ovaries were pre-hybridized 200 μl hybridization buffer (300 mM NaCl, 30 mM Sodium citrate pH 7.0, 15 v/v % ethylene carbonate, 1 mM EDTA, 50 μg/ml heparin, 100 μg/ml salmon sperm DNA, 1 v/v % Triton-X-100) for 10 minutes at 42°C. All probes were labelled as described previously (Gáspár et al. 2017). 50 μl of pre-warmed probe mixture (12.5-25 nM/individual oligonucleotide) was added and hybridization was allowed to proceed for 2 hours at 42°C. Samples were washed by pre-warmed hybridization buffer, hybridization buffer: PBSTX 1:1 mixture, PBSTX, each for 10 minute at 42°C. Finally, pre-warmed PBSTX was added and the sample was allowed to cool down to room temperature. Ovaries were mounted in 80 v/v % 2,2-thiodiethanol in PBS. Stacks of images were acquired on a Leica TCS SP8 confocal microscope using a 63x 1.4 NA oil immersion objective. Images were processed using ImageJ (http://rsb.info.nih.gov/ij/). Granularity (area covered by *oskar* mRNA positive sponge bodies / total oocyte area) was used as a quantitative measure for reticulated sponge body formation.

### Immunostaining of egg chambers

For the visualization of acetylated α-tubulin, ovaries from 2-day-old well-fed females were dissected into ice cold methanol and fixed for 20 minutes at -20°C. Ovaries were subsequently washed and rehydrated with PBS for 1 hour, blocked in 10% goat serum in PBS for 1 hour and stained overnight at 4°C with mouse anti-acetylated-α-Tubulin antibody. After washing, ovaries were incubated with a secondary antibody donkey anti-mouse-Cy3 (Jackson ImmunoResearch 715-165-151) and stained with DAPI. For visualization of Me31B, dGe1 and DCP1, ovaries were dissected into 2% v/v PFA (Electron Microscopy Sciences, #15710), 0.05% v/v Triton X-100 (Sigma, P1379) in PBS (pH 7.4) and fixed for 20 minutes. After washes in PBST (0.1% Tween20 in PBS), ovaries were permeabilized in 1% Triton X-100 in PBS for 1 hour, blocked with 10% goat serum in PBS and stained with primary antibodies in blocking buffer. After washing, ovaries were incubated with a secondary antibody, donkey anti-mouse-Cy3 (Jackson ImmunoResearch, 715-165-151) for Me31B and dGe1, and anti-GFP antibody-Alexa Fluor 488 conjugate (Invitrogen, A-21311) for YFP-DCP1. The samples were then stained with DAPI.

Stacks of images were acquired on a Leica TCS SP8 LIGHTNING confocal microscope using a 63x 1.4 NA oil immersion objective. Images were processed and analysed using ImageJ (http://rsb.info.nih.gov/ij/).

### Immunostaining of S2R+ cells

HDAC1 was amplified from *w*^*1118*^ cDNA using the primer pair CACCATGCAGTCTCACAGCAAAAAGC/TCAAATGTTGTTCTCCTTGGCG and inserted into pENTR/D-TOPO (Invitrogen). S391A and S391D mutations were introduced by site directed mutagenesis. The inserts were then cloned into the destination vector pAGW (The Drosophila Gateway Vector Collection) using the Gateway LR Clonase II (Thermo, 11791100). pA-GFP-HDAC1 wt/S391A/S391D were transfected into S2R^+^ cells using Effectene (Qiagen, 301425). 24 hours post transfection, cells were fixed with ice cold methanol for 15 minutes at -20°C. After permeabilized with 0.1% Triton X-100, cells were blocked with 10% goat serum in PBS and stained with mouse acetylated-α-Tubulin. After washing, cells were incubated with a secondary antibody donkey anti-mouse-Cy3 (Jackson ImmunoResearch 715-165-151) and stained with DAPI. Stacks of images were acquired on a Leica TCS SP8 LIGHTNING confocal microscope using a 63x 1.4 NA oil immersion objective. Images were processed and analysed using ImageJ (http://rsb.info.nih.gov/ij/).

### *Ex vivo* culturing of egg chambers

Ovaries from *w*^*1118*^ females kept on yeast paste were extracted and kept in Schneider’s *Drosophila* Medium (Gibco, 21720-024) supplemented with 10% FBS for 2.5 hours at room temperature. For the inhibition of phosphatases, the medium was supplemented with PhosSTOP (Roche, 4906845001), for the inhibition of CK2 the medium was supplemented with indicated amounts of TBCA (Sigma, SML0854). DMSO was used as control. Ovaries were harvested and subjected to analysis by western blot or fluorescent in situ hybridization.

### Deacetylation of tubulin and short peptides

*Drosophila* HDAC1 (wt/S391A/S391D) was cloned into pCoofy41 using SLIC cloning (Scholz et al. 2013). HDAC-TwinStrep was expressed in Sf21 cells using a baculovirus-mediated expression. Cell pellets were resuspended in Lysis Buffer (50 mM Tris, 150 mM NaCl, 1mM EDTA, 0.01% NP40, cOmplete Mini EDTA-free Protease inhibitor cocktail (Roche) and Benzonase) and lysed in a Microfluidizer (M-110L, Microfluidics). The lysate was cleared by centrifugation at 40,000 × g for 30 min at 4 °C. The supernatant was loaded onto a StrepTrap HP column (GE Healthcare), washed with Binding Buffer (50 mM Tris, 150 mM NaCl, 1mM EDTA) and eluted with 2.5 μM desthiobiotin in Binding Buffer. The eluted protein was supplemented with 5% glycerol. Single use aliquots were flash frozen in liquid nitrogen and stored at -80°C. For assaying the α-tubulin deacetylase activity of recombinant HDAC1, 1 μg tubulin purified from porcine brain (Cytoskeleton, T240) was mixed with 0.5 μg/μl BSA in assay buffer (50 mM Tris, 150 mM NaCl, 1mM EDTA, 3 mM KCL, 1 mM MgCl, 10% Glycerol) and supplemented with indicated amounts of HDAC1-TwinStrep in a 20 μl reaction. After an incubation for 1 hour at 25°C, acetylation of α-tubulin was analysed by western blot, probing against acetylated α-tubulin. Acetylated α-tubulin levels were normalized to total α-tubulin.

For the peptide deacetylation assay, recombinant HDAC-TwinStrep wild type, S391A or S391D activity was measured using a fluorimetric activity assay (Fluor de Lys, Enzo Life Sciences, BML-AK500-0001) according to the manufacturer instructions.

### Quantitative phosphoproteomics of well-fed and nutrition deprived egg chambers

#### Sample preparation

Freshly eclosed Oregon-R flies were kept in cages for 1-3 days at 25°C on protein rich diet (yeast paste on apple juice-agar plates). To provoke the nutritional stress, yeast paste was removed after 3 days for 4.5 hours prior to the preparation of egg chambers. Flies were narcotized with CO_2_, passed through twice a grain mill (KitchenAid) with PBS and sieved through meshes with aperture sizes of 630 μm, 400 μm, 2 × 200 μm, and egg chambers were collected in a mesh with aperture sizes of 80 μm. Egg chambers were collected in 50 ml tube and washed two times with PBS. Each sample was lyzed in 200 mM HEPES buffer (pH 8) containing PhosSTOP phosphatase inhibitors (Roche), phosphatase inhibitor cocktail2 (Sigma), cOmplete Mini EDTA-free Protease inhibitor cocktail (Roche), and RapiGest SF surfactant (Waters) using a dounce homogenizer. The samples were incubated for 5 minutes at 70°C in a water bath and sonicated for 20 minutes in a water bath. After centrifugation for 5 minutes at 4’000 g, the supernatant was transferred to a new vial and the protein concentration was determined via a bicinchoninic acid (BCA) assay. The proteins were reduced with dithiothreitol (DTT) with subsequent carbamidomethylation of cysteine residues with iodoacetamide (IAA). The modified proteins were digested with lysyl endopeptidase (Lys-C) (Wako) for 3h at 37° C and subsequently digested with trypsin (Promega) overnight at 37° C. After the digestion, RapiGest SF surfactant was cleaved after addition of 10% trifluoroacetic acid (Biomol) during 45 min at 37°C. After centrifugation for 20 min at 4°C at 16,000 g, the supernatant was desalted on a SepPak 1 cc 50 mg cartridge (Waters). The eluate was concentrated in a vacuum centrifuge. After addition of HEPES buffer, the pH was adjusted to 7.7 with NaOH and/or HCl.

#### TMT labeling

The TMT-reagent (Thermo) was directly added to the sample and incubated for 1 hour at 23°C shaking at 300 rpm in a heating block. The reaction was stopped by addition of 8 μl of 5% hydroxylamine and incubation for 15 min at 23°C shaking at 300 rpm in a heating block. The sample was desalted again on a SepPak 1 cc 50 mg cartridge (Waters). The procedure was repeated in order to reach complete labelling. Prior to mixing all six samples, a test mixture was created and analysed by LC-MS. Based on the results of this analysis all TMT-labelled samples were mixed in a 1:1 ratio.

#### TiO_2_ phosphopeptide enrichment

For the mixed TMT labelled samples a microcolumn was packed. A plug of an Empore C8 disk (3M) was pushed into an GELoader tip (Eppendorf). The tip was further filled with Titansphere 5 μm TiO_2_ beads (GL Science). The microcolumn was washed twice with 6% trifluoroacetic acid (TFA) in 80% acetonitrile (MeCN). The flow through from the sample loading was collected for further analysis. Subsequently, the microcolumn was washed first with 6% TFA in 80% MeCN, second with 200 mM NaCl in 50% MeCN acidified with 0.5% TFA and third with 0.1% TFA in 80% MeCN. The sample was eluted with 5% ammonia and collected in 10% formic acid. A second elution with 0.1% TFA in 80% MeCN was collected in the same vial.

#### Prefractionation

The flow through and the enriched phosphopeptides were adjusted to a basic pH with 25% ammonium just before injection. The separation was performed on an Agilent 1260 infinity high-performance liquid chromatography (HPLC) system equipped with a Waters XBridge C18; 3,5 μm; 1 × 100 mm reversed phase column at a flow rate of 75 μl/min. The running buffers were 20 mM ammonium formate at pH 10 and 100% acetonitrile. In total, 90 one minute fractions were collected in a 96 well plate containing 25 μl 1% formic acid per well to neutralize the basic pH of the running buffer. The fractions were concentrated in a vacuum centrifuge and desalted and pooled in one step as reported previously (Hennrich et al. 2018).

#### NanoLC-MS/MS analysis

The resulting 18 fractions from the phosphopeptide enriched sample and the 18 fractions from the flow through sample were separated by an UltiMate 3000 RSLC nano LC system (Dionex). The peptides were trapped on a μ-Precolumn C18 PepMap 100 (300 μm × 5 mm, 5 μm, 100 Å) and separated on an Acclaim PepMap 100 C18 (75 μm × 50 cm, 3 μm, 100 Å) column. The analytical column was connected to a Pico-Tip Emitter (New Objective; 360 μm OD x 20 μm ID; 10 μm tip). The LC system was coupled to a QExactive plus (Thermo) via a proxeon nanoflow source. The solvents used for peptide separation were 0.1% formic acid in water and 0.1% formic acid in acetonitrile. The flow rate was 30 μl/min for trapping and 300 nl/min for the separation. The initial conditions were 2% of organic phase isocratic for 2.9 minutes followed by an increase to 4% in 1.1 minutes and 8% in 2 minutes. The subsequent linear gradient from 8% to 28% organic solvent was 96 minutes long and was followed by a 10-minute gradient from 28% to 40% organic solvent. The washing of the column was at 80% organic solvent for 2 minutes followed by re-equilibration at 2%. Full scan spectra were acquired in positive ion mode in a mass range of 375-1200 m/z in profile mode with the resolution set to 70,000, with an AGC target of 3×106 ions and a maximum ion trap fill time of 30 ms. Fragmentation spectra were recorded at a resolution of 35,000 in the orbitrap with an AGC target of 2×105 ions and a maximum ion trap fill time of 120 ms. A top10 method was chosen with a precursor isolation window of 0.7 m/z and a fixed first mass of 100 m/z. The normalized collision energy was set to 32 and spectra were recorded in profile mode. Unassigned peaks as well as charge states 1, 5-8 and >8 were not chosen for fragmentation. The peptide match algorithm was set to ‘preferred’ and the dynamic exclusion to 30 seconds.

#### Data processing

The raw data files were processed with Thermo Proteome Discoverer version 1.4.1.14. The mass range from 126-131.3 m/z was excluded in the fragmentation spectra. In addition, the spectra were de-isotoped, deconvoluted and a top N filter was applied with N set to 10 in a 100 Da window prior to database search with Mascot version 2.5. The spectra were searched against the Uniprot database *Drosophila melanogaster* including common contaminants (22157 sequences). Semitrypsin was chosen as enzyme with one missed cleavage allowed. Further settings were a precursor mass tolerance of 10 ppm, a fragment mass tolerance of 0.02 Da, carbamidomethylation set as fixed modification, oxidation on methionine and phosphorylation on serine, threonine and tyrosine set as variable modification. The quantification method included TMT as fixed modification. Percolator version 2.04 was used for false discovery rate (FDR) calculation and PhosphoRS 3.0 for calculating site probabilities for all potential phosphorylation sites. The results were exported, normalized with the vsn method (Huber et al. 2002) and further processed by a custom analysis pipeline based on the Limma package in R/Bioconductor (Ritchie et al. 2015). Protein fold-changes were calculated from flow through samples and used for normalization of phospho-site fold-changes. GO term analysis was performed using the FlyMine database (Lyne et al. 2007).

## Data availability

The mass spectrometry phosphoproteomics data have been deposited at the Open Science Framework repository ([OSF](https://osf.io/dprt5/?view_only=b3770945c8384fbe962da1518e4f84be)).

## Code availability

The computer code used for analysis of the phosphoproteomics data is available from: https://github.com/frankwippich/Wippich_Phosphoproteomics-analysis

## Acknowledgements

We thank Jürg Müller (Max Planck Institute of Biochemistry, Martinsried, Germany) for the anti-HDAC1 antibody. We are grateful to the members of the Ephrussi laboratory for helpful discussions and critical reading of the manuscript. F.W. was supported by a postdoctoral fellowship from the EMBL Interdisciplinary Postdoc Program (EIPOD) under Marie Curie COFUND actions (European Union). This work, F.W. and V. were supported by DFG-FOR 2333 grants EP 37/2-1 and and EP 37/4-1 from the Deutsche Forschungsgemeinschaft (Germany) to A.E. and by the EMBL.

## Author Contributions

F.W., V. and A.E. designed the study, discussed the results and wrote the manuscript. F.W. and V. designed, performed and analysed the experiments. F.W. and M.H. designed, conducted and analysed the quantitative phosphoproteomic assay.

---

**Supplemental Figure S1:**
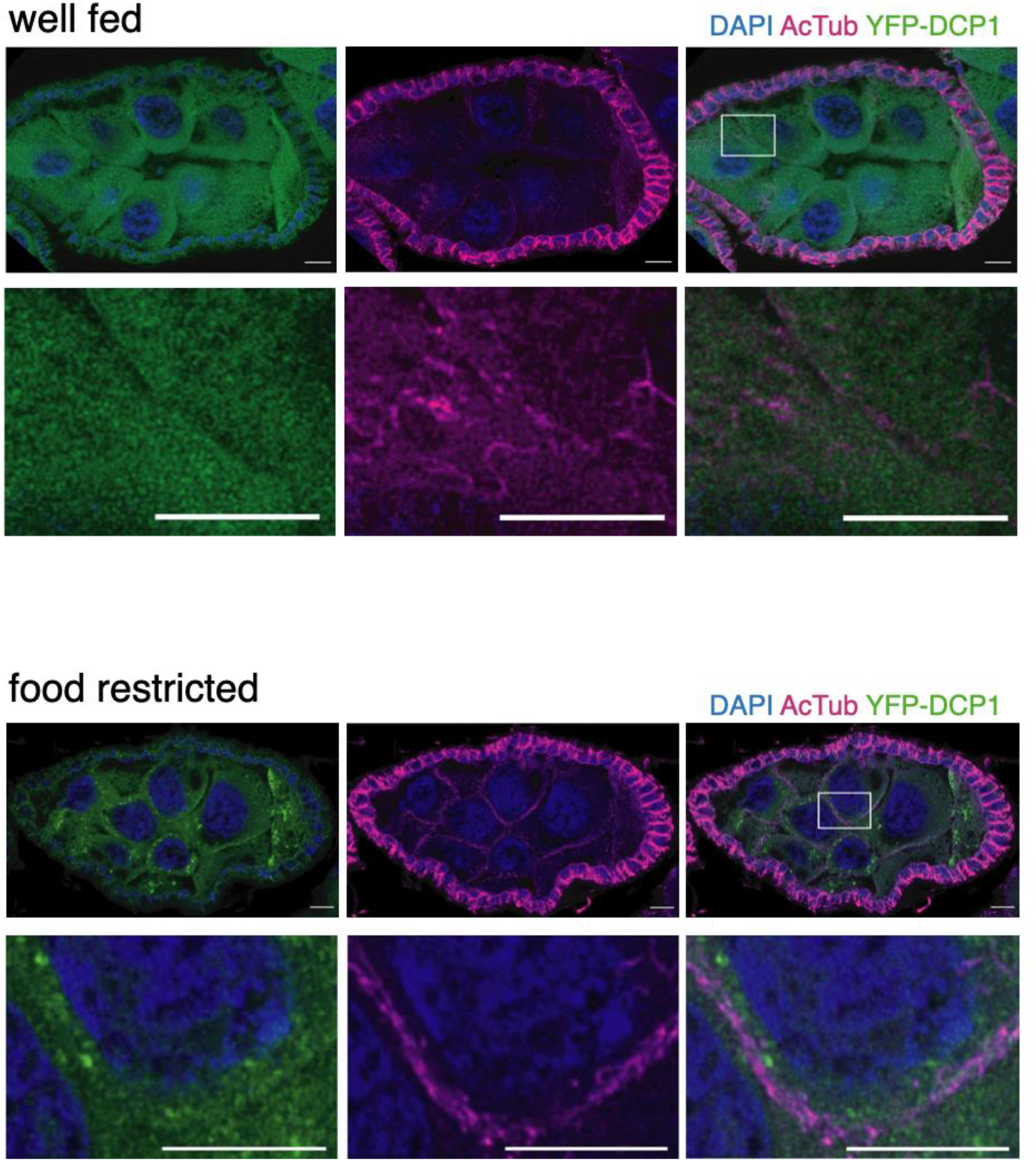
Starvation induces sponge body formation and cortical accumulation of acetylated microtubules. Sponge bodies are visualized by staining for YFP-DCP1, a known sponge body marker (Lin *et al*, 2006). The lower panels show magnified images of a cell boundary within the rectangular area highlighted in the upper panels on the right (merged signals). Scale bar: 10μm

**Supplemental Figure S2:**
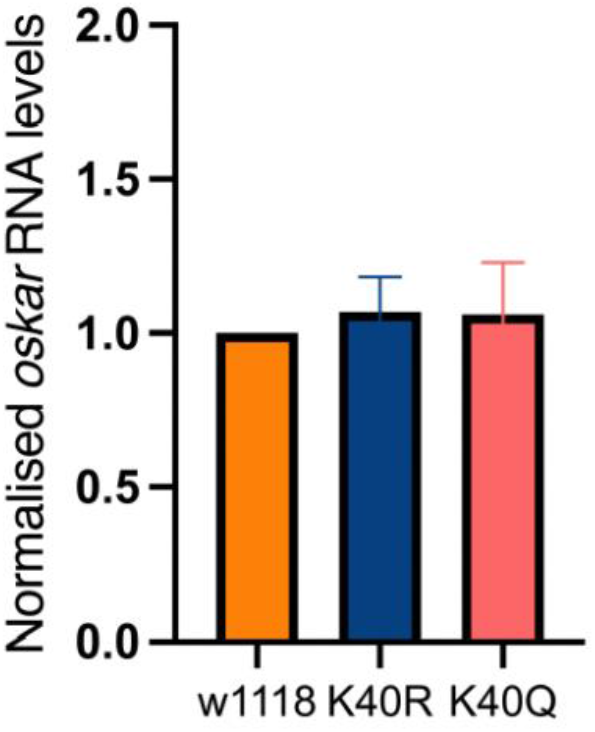
*oskar* mRNA levels are unaffected in K40R, K40Q expressing flies compared with the w1118 control. *oskar* mRNA levels were quantified by qPCR and normalized to total 18S rRNA levels.

**Supplemental Figure S3:**
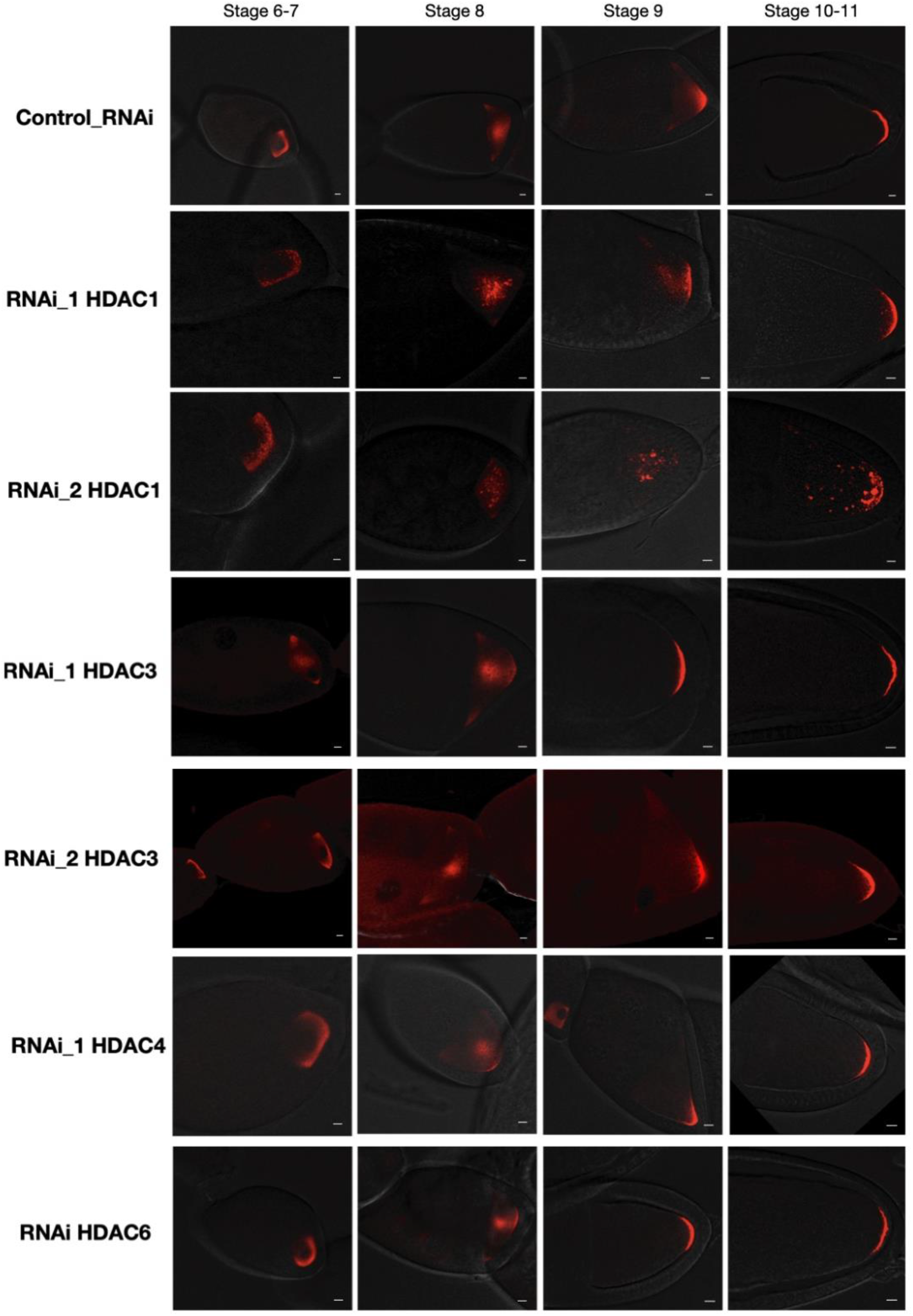
Knock-down of HDAC1, but not other histone deacetylases, leads to mRNA aggregation in well-fed flies. *oskar* mRNA detected by smFISH. Panels from left to right show oocytes at progressive stages of development. Scale bars: 5μm

**Supplemental Figure S4:**
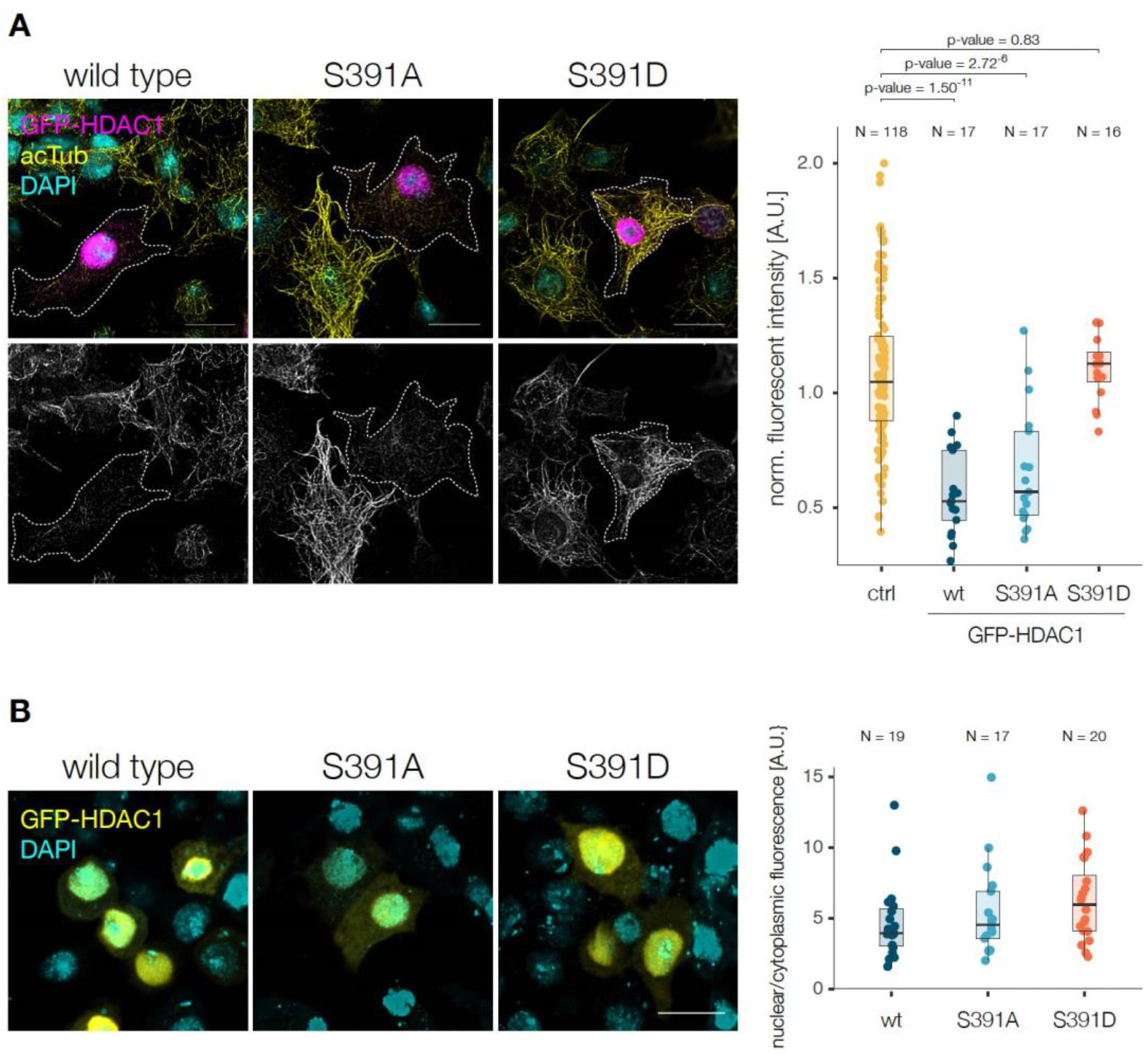
HDAC1 wild type and mutant overexpression in S2R+ cells. (A) S2R+ cells transiently transfected with GFP-HDAC1 wild type, S391A or S391D stained 24 hours post transfection for acetylated α-tubulin K40 and DAPI. The fluorescence intensity of acetylated α-tubulin K40 was quantified. Non transfected cells served as control. Statistical significance was calculated using Welch Two Sample t-test. (B) The nucleocytoplasmic ratio of S2R+ cells overexpressing GFP-HDAC1 wild type, S391A or S391D was quantified. Scale bars: 10 μm.

**Supplemental Figure S5:**
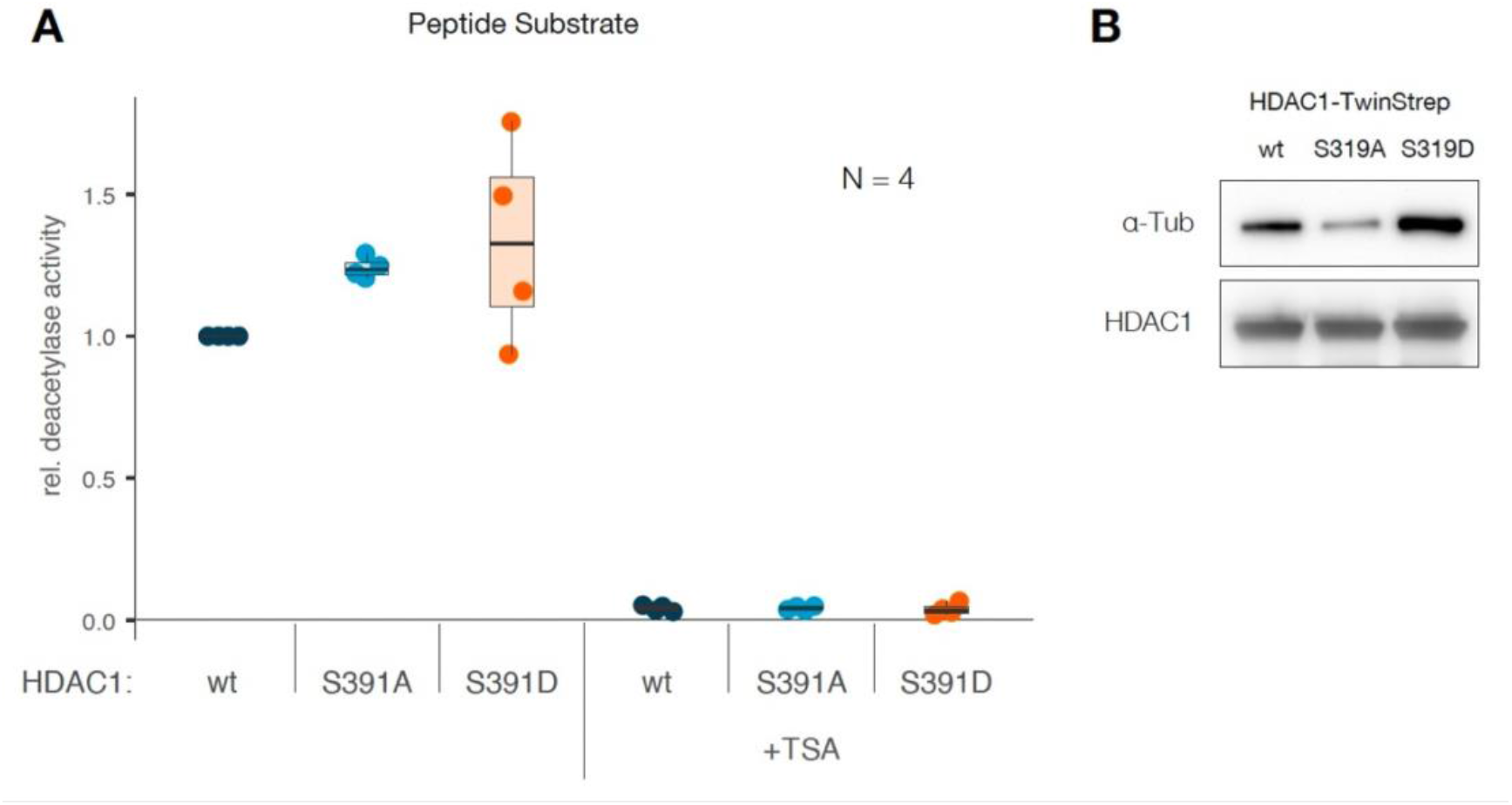
The effect of HDAC1 S319 mutations on the deacetylation activity of short peptide substrates and α-tubulin binding. (A) Recombinant HDAC1 wild type, S391A or S391D was used in a fluorimetric activity assay using short peptide substrates. The error bars represent standard deviation. (B) Western blot analysis of α-tubulin copurified with HDAC1-TwinStrep wild type, S391A or S391D expressed in insect cells.

**Supplemental Figure S6:**
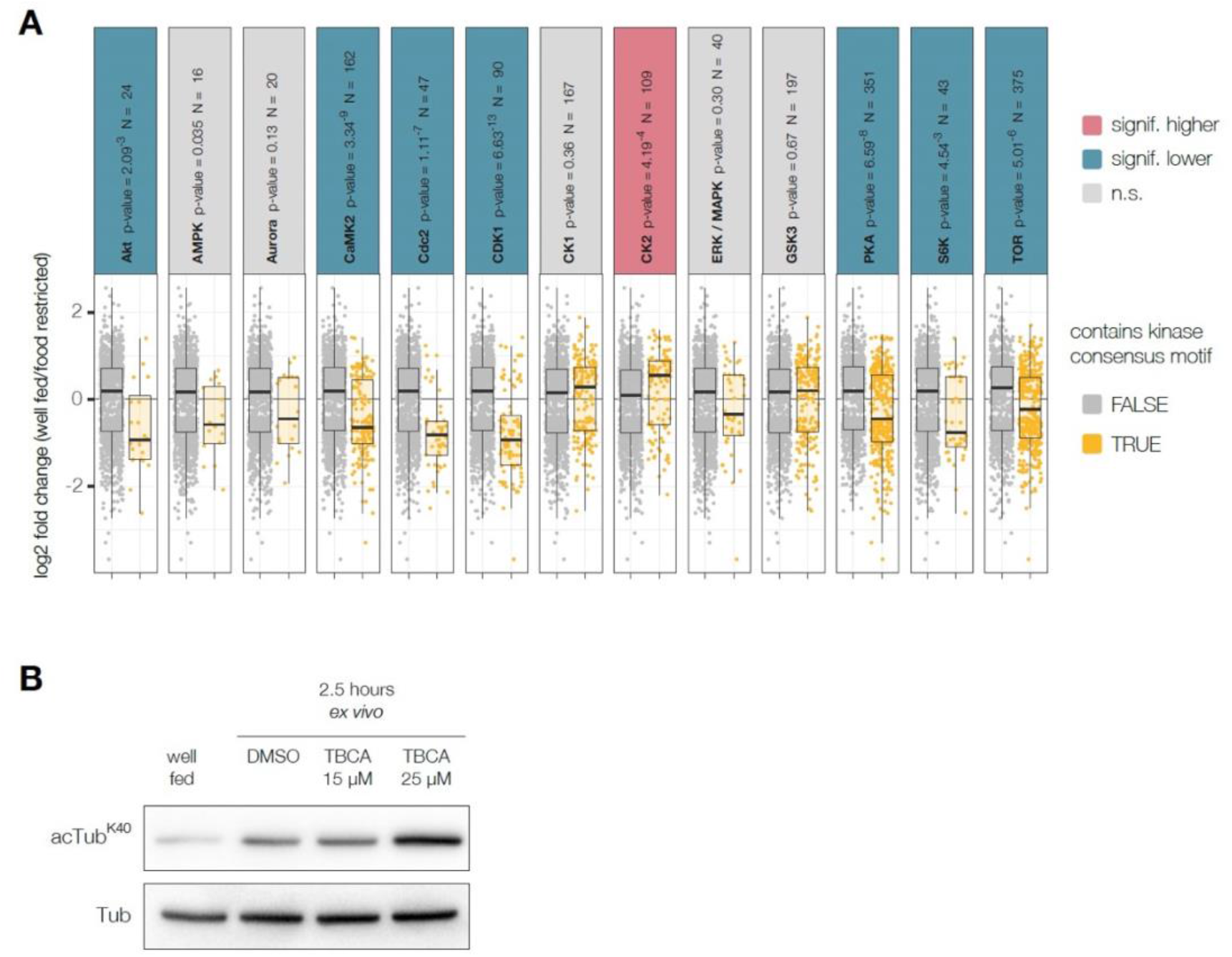
Kinases involved in nutritional stress transmission. (A) Analysis of the response of phosphorylation sites belonging to the indicated kinase consensus motif group to nutritional stress. Kinase consensus motif groups that change significantly are highlighted (blue: less phosphorylated during nutritional stress; red: more phosphorylated during nutritional stress). Statistical significance was calculated using Wilcoxon rank sum test. (B) Acetylation of α-tubulin in ovaries from well fed females or *ex vivo* cultured in the presence of DMSO control or TBCA was analysed by western blot.

**Supplemental Figure S7:**
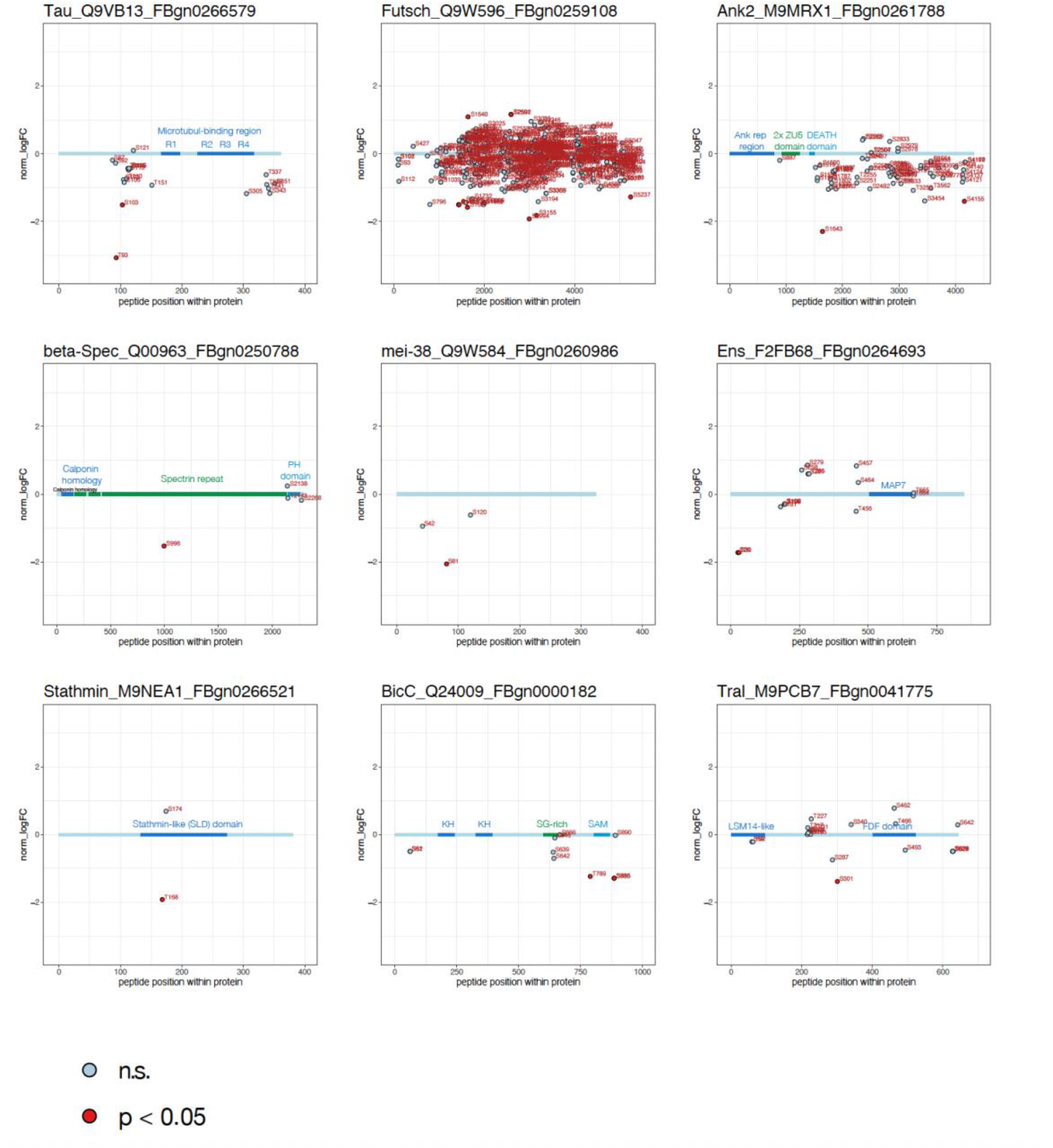
Examples of microtubule associated proteins whose phosphorylation status was significantly altered in response to nutritional stress. Visualization of example proteins (Name_UniProtID_FlybaseID), domains and the phosphorylation sites identified by quantitative phosphoproteomics.

